# Sleep regulates hepatic neutrophil trafficking by modulating uric acid metabolism via sympathetic nervous system activity

**DOI:** 10.64898/2026.04.28.721287

**Authors:** Pawan K. Jha, Utham K. Valekunja, Jing Chen-Roetling, Akhilesh B. Reddy

## Abstract

**Sleep is essential for survival and serves as a key regulator of metabolic and immune function. Sleep loss is strongly associated with metabolic stress and liver inflammation. The mechanisms linking sleep disruption to hepatic metabolic inflammation (metaflammation) remain poorly defined. Here, we show that sleep loss triggers metaflammation through a sympathetic-metabolic-immune axis. Acute sleep deprivation (SD) activates hepatic sympathetic signaling, leading to increased uric acid (UA) synthesis driven by enhanced expression and activity of xanthine dehydrogenase/xanthine oxidase (XDH/XO) in the liver. Elevated UA, acting as an immune-stimulatory metabolic signal, promotes hepatic neutrophil recruitment and pro-inflammatory cytokine production, a response that is rapidly reversed upon sleep recovery. Our findings identify sleep-dependent sympathetic control of hepatic UA metabolism as a driver of acute liver inflammation and reveal how acute sleep loss reprograms liver immune-metabolic homeostasis.**

## Introduction

Metabolic homeostasis is sustained by dynamic crosstalk between the brain and peripheral tissues, in which central neural circuits regulate metabolism. Within this network, the liver serves as a central regulator of glucose and lipid homeostasis ^1^. Sleep is a critical physiological process required to maintain this metabolic homeostasis. Sleep disruption is strongly associated with metabolic dysregulation or increased susceptibility to metabolic diseases ^2–6^. Sleep deprivation has been reported to induce liver inflammation and generate reactive oxygen species in hepatocytes ^7–9^. Emerging evidence suggests the existence of circuits connecting the sympathetic and immune systems through which the sleeping brain regulates metabolic inflammation. For instance, chronic sleep loss induces hepatic steatosis in mice through sympathetic nerve sprouting and aseptic inflammation ^10^. It remains unclear, however, how sleep-dependent sympathetic signaling to the liver controls immune cell trafficking and activity, thereby influencing metabolic inflammation.

Neutrophils are a class of innate immune cells that serve as first responders to inflammation. They rhythmically infiltrate the liver and regulate circadian gene expression to coordinate daily hepatic metabolism ^11^. Neutrophils also exhibit diurnal oscillations in their recruitment and migration across multiple tissues ^12^, and have been identified as key contributors to hepatic steatosis ^13^. Notably, hepatic neutrophil abundance is reduced during rest and increased during the active phase ^11^, suggesting that sleep could actively gate neutrophil trafficking to the liver. Prolonged sleep deprivation has been shown to induce liver tissue necrosis and inflammatory cell infiltration ^14, 15^, yet the mechanisms linking sleep loss to dysregulated hepatic immune cell dynamics are not defined.

Here, we aim to address this knowledge gap by elucidating the mechanisms through which sleep regulates hepatic neutrophil trafficking and, in turn, liver function. We describe a sleep-dependent circuit that regulates neutrophil trafficking and inflammatory responses. We identified that sleep deprivation activates the sympathetic branch of the autonomic nervous system, which innervates the liver, leading to increased hepatic uric acid (UA) synthesis. Elevated UA, a damage-associated molecular pattern (DAMP), promotes neutrophil influx into the liver and the production of pro-inflammatory cytokines. Together, these findings reveal a mechanism by which sleep modulates inflammatory mediators to maintain inflammatory homeostasis via sympathetic tone. Furthermore, our results highlight the therapeutic potential of targeting this axis to rebalance metabolic inflammation induced by sleep loss.

## Results

### Sleep regulates liver neutrophil recruitment and sterile inflammation

We first investigated whether acute loss of sleep affects neutrophil recruitment to the liver. Mice were assigned to three experimental groups: a normal sleep (NS) control group, a 12-h (ZT0 to ZT12, ZT; zeitgeber time) sleep-deprived (SD) group, and a sleep-deprivation followed by 24-h recovery sleep (RS) ^16^ (Fig. 1a). To control for circadian influences on neutrophil distribution, liver and blood samples were collected from all groups at the same circadian time point (ZT12) (Fig. 1a). We then assessed neutrophil influx and activation status in the liver. We found a marked increase in neutrophil recruitment and in the proportion of activated neutrophils, as indicated by myeloperoxidase (MPO□) expression, in the livers of SD mice. These changes reverted to NS levels following a sleep recovery (Fig. 1b, Supplementary Fig. 1a-b). These hepatic alterations were paralleled by corresponding changes in circulating neutrophil proportions (Supplementary Fig. 2a-e), suggesting enhanced neutrophil migration to the liver. In agreement with previous reports ^14^, we also saw a reduction trend in circulating lymphocyte proportions following SD (Supplementary Fig. 2c). Histopathological analyses further confirmed the increased hepatic infiltration of neutrophils in an activated state (Fig. 1c-d).

**Fig 1.**
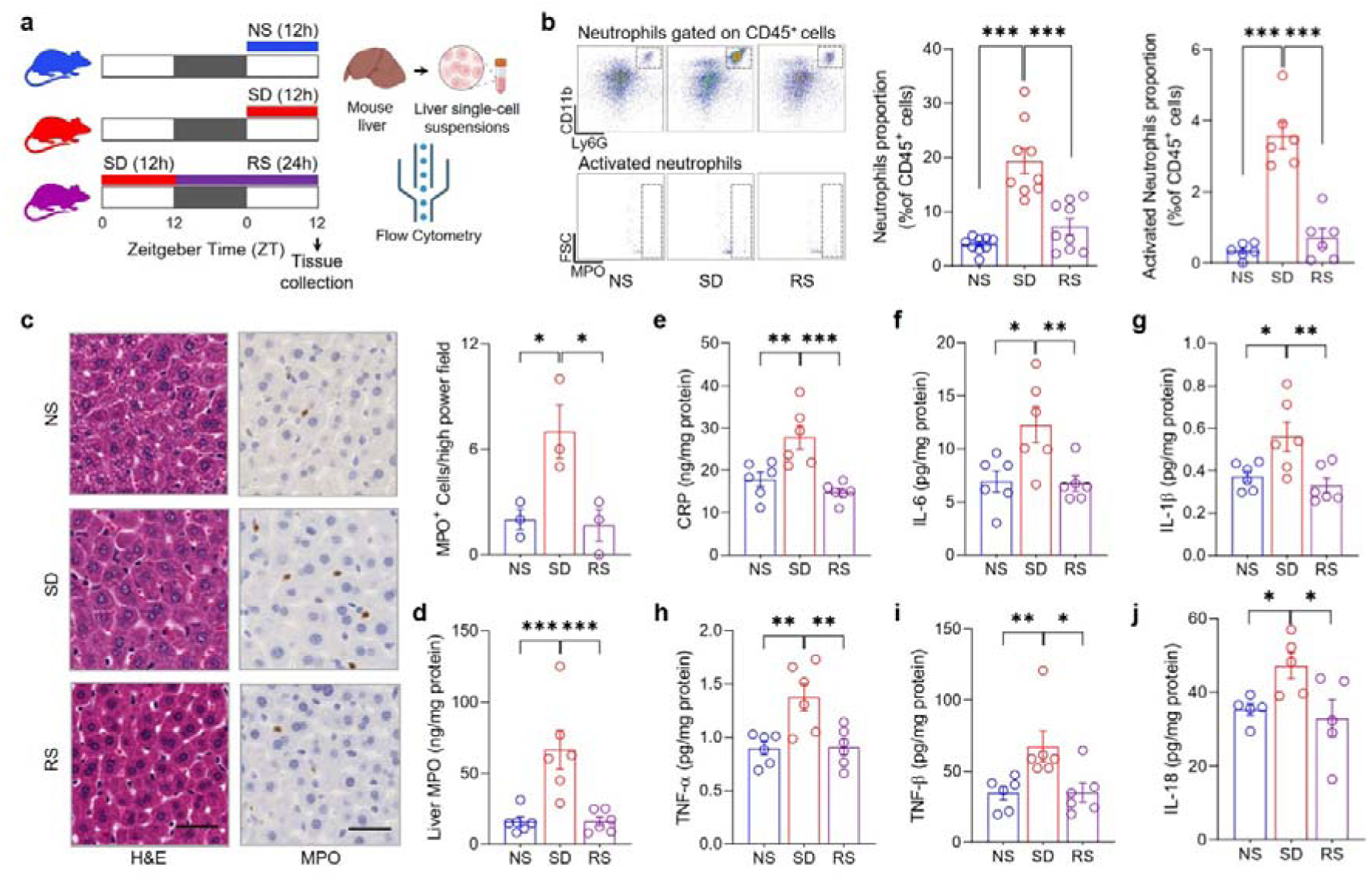
Sleep regulates hepatic neutrophil influx and aseptic inflammation. **a,** Schematic of experiment design for sleep treatments, tissue collection, and preparation of liver single-cell suspension for flow cytometry (black bars represent darkness, white bars indicate periods of light). **b,** Gating strategy and bar plots showing quantification for neutrophils (upper, left), activated neutrophils (lower, right). **c,** Representative H&E (arrow indicates granulocyte), MPO-stained sections, and MPO +ve cells (brown) quantification (bar plot, right) of liver from sleep treatment groups. Bar 40 µm. **d-j,** Bar plots showing alterations of liver MPO (d), CRP (e), IL-6 (f), IL-1β (g), TNF-α (h), TNF-β (i), and IL-18 (j) across sleep treatments. Data are mean□±□s.e.m. n□=□3-9 biological replicates. One-way ANOVA followed by post-hoc Fisher’s Least Significant Difference (LSD) test. \**P* < 0.05, \*\**P* < 0.01, and \*\*\**P* < 0.001. NS normal sleep, SD sleep deprived, RS recovery sleep, H&E hematoxylin & eosin, MPO myeloperoxidase, CRP C-reactive protein, IL Interleukin, TNF tumor necrosis factor.

To map out the inflammatory environment linked to the influx of neutrophils into the liver associated with SD, we measured several important inflammatory signals. Most proinflammatory markers were sharply elevated in the livers of SD mice and returned close to normal levels after 24 h of recovery sleep (Fig. 1e-j, Supplementary Fig. 1c-e, Supplementary Fig. 3h-j). In contrast, analysis of the systemic response in the plasma revealed that only CRP, IL-6, IL-1β, and TNF-α exhibited a similar pattern of elevation after SD (Supplementary Fig. 2f-n). Together, these results suggest that acute sleep loss preferentially induces a localized inflammatory response in the liver rather than a broad systemic inflammatory response.

Sleep disruption induces hepatic metabolic stress ^17, 18^. Accordingly, we assessed whether metabolic perturbations in the liver coincided with inflammatory responses. We measured hepatic mRNA expression of genes governing metabolism and liver function (*G6pc*, *Pck1*, *Sirt1*, *Ppar*α, *Nr1d1*, and *Ho-1*) in mice subjected to the indicated sleep interventions (Supplementary Fig. 3a-g). Strikingly, the temporal pattern and reversibility of these metabolic changes closely tracked neutrophil recruitment and activation in the liver, supporting a tight coupling between acute metabolic stress and neutrophil-driven inflammation.

### Sleep deprivation increases hepatic UA synthesis

We next sought to identify sleep-dependent metabolic signals in the liver that drive neutrophil recruitment. UA is the final breakdown product of purine metabolism and is mainly produced in the liver. It acts as a damage-associated molecular pattern (DAMP), meaning it can trigger inflammatory responses driven by neutrophils ^19–21^. We therefore asked whether acute sleep deprivation alters hepatic UA biosynthesis. We profiled metabolites in the UA synthesis pathway in the livers of mice subjected to sleep interventions. Hepatic UA was significantly elevated during SD and returned to baseline following recovery sleep (Fig. 2a). While most upstream metabolites remained unchanged, levels of inosine monophosphate (IMP), a central intermediate in purine metabolism, remained elevated during SD and persisted after 24 h of recovery. These findings indicate that SD could preferentially affect the terminal steps of UA production, namely the conversion of hypoxanthine and xanthine to UA.

**Fig 2.**
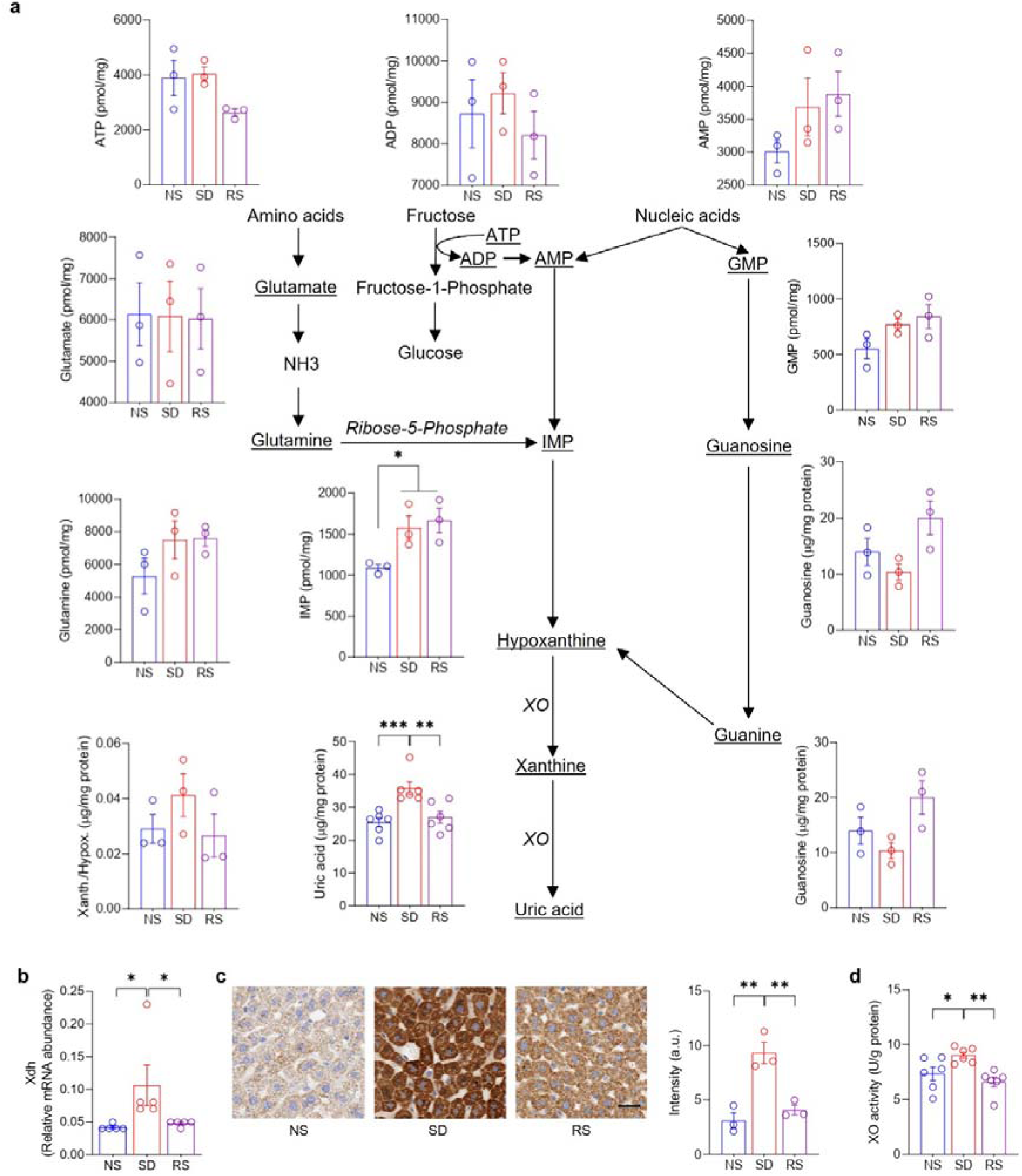
Sleep deprivation increases hepatic uric acid (UA) levels through the actions of Xanthine dehydrogenase (Xdh)/Xanthine oxidase (Xo). **a,** Bar plots showing targeted metabolomics and fluorometric quantification of uric acid pathway metabolites across sleep treatments. **b**, mRNA expression of *Xdh* normalized to *Actb*. **c,** Representative XO-stained sections, and intensity quantification (bar plot, right) of liver from sleep treatment groups. Bar 30 µm. **d**, Bar plot showing XO activity from the liver lysate of animals subjected to different sleep treatments. Data are mean□±□s.e.m. n□=□3-6 biological replicates. One-way ANOVA followed by post-hoc Fisher’s Least Significant Difference (LSD) test. \**P* < 0.05, \*\**P* < 0.01, and \*\*\**P* < 0.001. NS normal sleep, SD sleep deprived, RS recovery sleep.

We therefore hypothesized that SD enhances the activity of xanthine oxidase/dehydrogenase (XO/XDH), the enzymes catalyzing this final step of UA synthesis, encoded by the *Xdh* gene. Consistent with this hypothesis, hepatic *Xdh* mRNA expression, XO enzymatic activity, and XO protein levels were all increased during SD and normalized after sleep recovery (Fig. 2b-d). By contrast, neither UA levels nor XO activity in the systemic circulation were altered by sleep manipulation (Supplementary Fig. 4a-b), indicating a liver-restricted effect. These data demonstrate that acute sleep loss selectively augments hepatic UA synthesis through enhanced XO/XDH expression and activity.

### Inhibition of UA synthesis attenuates SD-driven neutrophil influx and inflammation

Metabolic stress and tissue insult can promote UA generation from purine metabolism via XO, thereby acting as DAMPs that recruit neutrophils and trigger sterile inflammation ^20^. We therefore tested whether pharmacological inhibition of UA synthesis modulates SD-induced hepatic inflammation (Fig. 3a-l, Supplementary Fig. 5a-m). We treated mice with allopurinol, a selective XO inhibitor, prior to SD. This treatment significantly lowered UA levels in both the liver and the blood (Fig. 3c, Supplementary Fig. 5e). We then assessed neutrophil recruitment and activation in the liver. In vehicle-treated mice, SD induced a robust increase in hepatic neutrophil infiltration and a coordinate increase in the proportion of activated neutrophils. Allopurinol treatment, however, markedly attenuated both neutrophil accumulation and activation in response to SD (Fig. 3a-b). We further examined whether allopurinol affected circulating white blood cell populations and observed no significant changes attributable to allopurinol treatment. Consistent with the previous experiment (Supplementary Fig. 2a-e), SD was associated with a reduction in circulating lymphocytes and a concomitant increase in neutrophils (Supplementary Fig. 5a-d).

**Fig 3.**
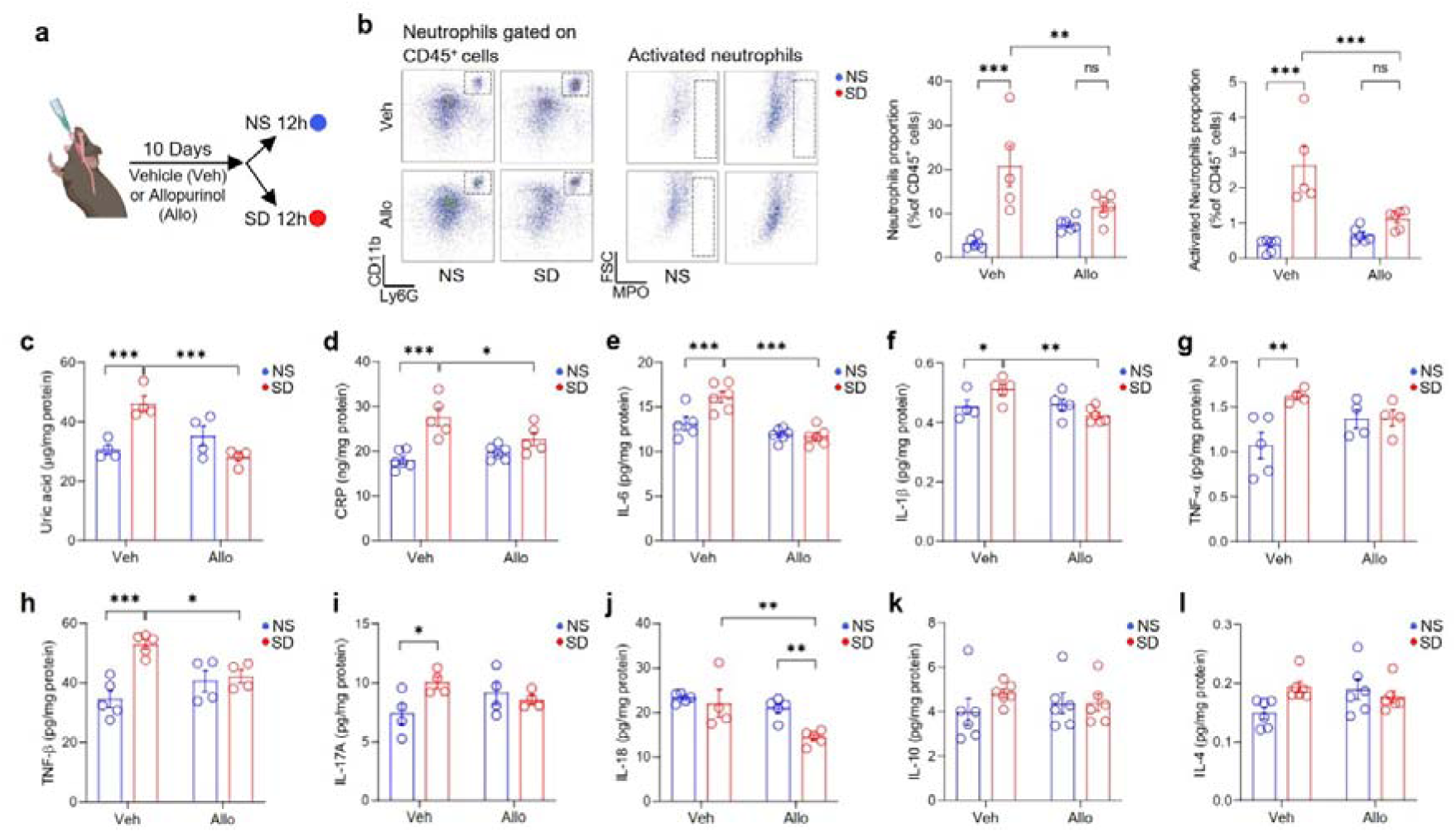
Inhibition of uric acid synthesis reduces hepatic neutrophil trafficking and inflammation following sleep deprivation. **a,** Schematic of allopurinol oral gavage and sleep deprivation. **b,** Gating strategy and bar plots showing quantification for neutrophils (left) and activated neutrophils (right) in the liver of mice treated with Allo/Veh followed by sleep treatments. **d-n**, Bar plots showing liver uric acid (c), CPR (d), IL-6 (e), IL-1β (f), TNF-α (g), TNF-β (h), IL-17A (i), IL-18 (j), IL-10 (k), and IL-4 (l) levels in the mice treated with Allo/Veh followed by sleep treatments. Data are mean□±□s.e.m. n□=□4-6 biological replicates. Two-way ANOVA followed by post-hoc Fisher’s Least Significant Difference (LSD) test. \**P* < 0.05, \*\**P* < 0.01, and \*\*\**P* < 0.001. NS normal sleep, SD sleep deprived, Veh vehicle, Allo allopurinol, UA uric acid, MSU monosodium urate, PBMC peripheral blood mononuclear cell.

We next asked whether the allopurinol-mediated reduction in hepatic neutrophil infiltration under SD conditions was accompanied by a corresponding attenuation of inflammatory signaling in the liver. Indeed, most proinflammatory mediators and cytokines were elevated during SD, and these responses were significantly blunted by allopurinol treatment (Fig. 3d-l). In contrast, analysis of circulating cytokines revealed a more limited effect, with only IL-6 showing a comparable reduction (Supplementary Fig. 5f-m), indicating that inhibition of UA synthesis primarily suppresses a liver-restricted inflammatory program induced by sleep deprivation.

### UA rise enhances neutrophil recruitment to hepatocytes

Neutrophil chemotaxis is a central feature of liver inflammation and metabolic stress ^22, 23^. Crystalline forms of UA, such as monosodium urate (MSU), can trigger such chemotactic responses ^20^. To determine whether MSU-stimulated hepatocytes generate signals that drive neutrophil migration, we performed transwell migration assays. MSU alone was insufficient to induce neutrophil migration directly. However, conditioned media from MSU-treated hepatocytes robustly promoted the migration of peripheral neutrophils from circulating WBCs (Fig. 4a). These findings suggest that elevated UA facilitates neutrophil recruitment indirectly by inducing hepatocytes to release soluble chemotactic factors.

**Fig 4.**
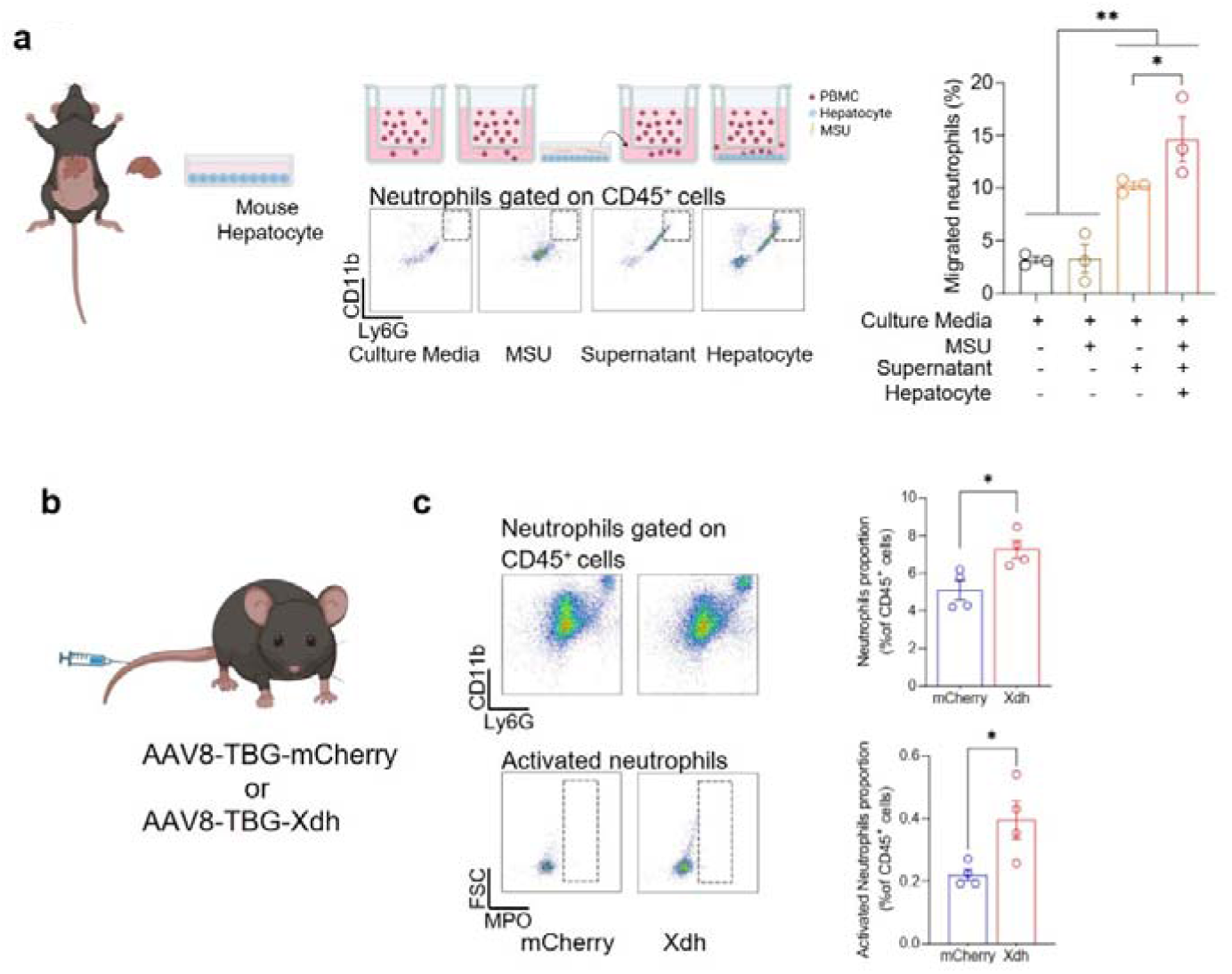
Uric acid crystals recruit neutrophils to hepatocytes, and liver-specific Xdh overexpression increases hepatic neutrophil infiltration. **a,** Experimental design and flow cytometry results of cultured mouse WBCs incubated with MSU or stimulated supernatant by Transwell assay. Bar plots showing migrated neutrophils. **b,** Schematic of liver-specific overexpression of mCherry and Xdh achieved by intravenous injection of AAV8-TBG-mCherry/Xdh. **c,** Gating strategy and bar plots showing quantification for neutrophils (upper) and activated neutrophils (lower) in the liver of mCherry or Xdh overexpressing mice. Data are mean□±□s.e.m. n□=3-4 biological replicates. One-way ANOVA followed by post-hoc Fisher’s Least Significant Difference (LSD) test and Unpaired t-test, two-tailed. *\*P* < 0.05, *\*\*P* < 0.01.

To determine whether this mechanism operates *in vivo*, we overexpressed *Xdh* specifically in hepatocytes to enhance endogenous UA production. *Xdh* overexpression resulted in a marked increase in hepatic neutrophil infiltration and a higher proportion of activated neutrophils (Fig. 4b-c). Notably, this phenotype closely recapitulated the neutrophil recruitment and activation seen during SD, supporting a causal role for UA-driven signaling in promoting hepatic neutrophil influx.

### Sympathectomy attenuates SD-induced hepatic neutrophil influx

SD is associated with heightened sympathetic nervous system (SNS) activity ^24, 25^, and stimulation of hepatic sympathetic nerves has been shown to promote UA production ^26^. We thus hypothesized that the SD-induced increase in hepatic UA levels and neutrophil recruitment is driven by enhanced sympathetic signaling. To test this, we performed global sympathectomy by intraperitoneal administration of 6-hydroxydopamine hydrochloride (6-OHDA) before SD challenges (Fig. 5a). Remarkably, sympathectomy evidently abrogated SD-induced neutrophil accumulation in the liver and significantly reduced the proportion of activated neutrophils (Fig. 5a–b), implicating SNS activity in regulating hepatic neutrophil dynamics during SD.

**Fig 5.**
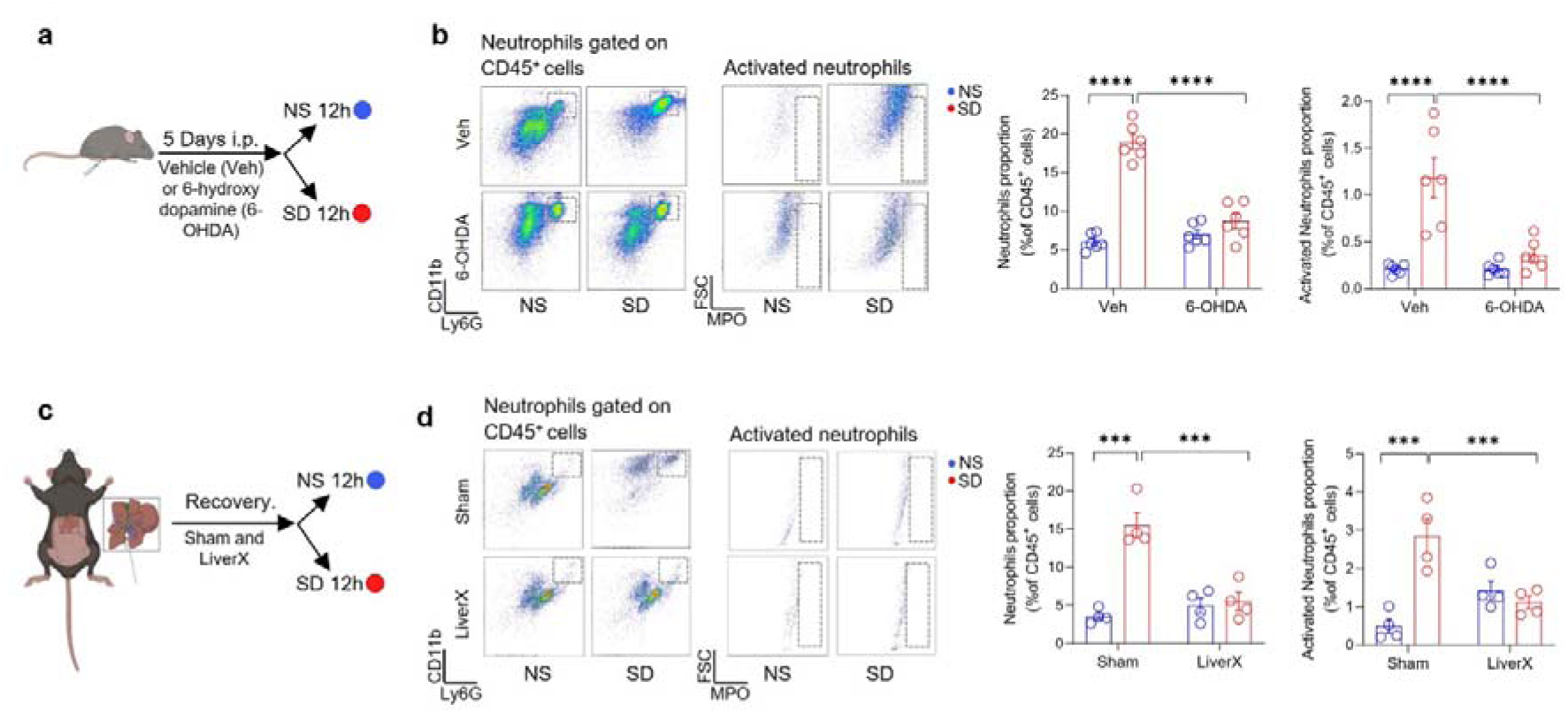
Both global and liver-specific sympathectomy reduce hepatic neutrophil infiltration following sleep deprivation. **a,** Schematic of 6-OHDA treatments followed by sleep deprivation. **b,** Gating strategy and bar plots showing quantification for neutrophils (left) and activated neutrophils (right) in the liver of mice treated with 6-OHDA followed by sleep treatments. **c**, Schematic of phenol-based liver sympathectomy followed by sleep deprivation. **d,** Gating strategy and bar plots showing quantification for neutrophils (left) and activated neutrophils (right) in the liver of mice that underwent hepatic sympathectomy followed by sleep treatments. Data are mean□±□s.e.m. n□=□4-6 biological replicates. Two-way ANOVA followed by post-hoc Fisher’s Least Significant Difference (LSD) test. *\*P* < 0.05, *\*\*P* < 0.01, and *\*\*\*P* < 0.001. NS normal sleep, SD sleep deprived, 6-OHDA 6-hydroxydopamine, LiverX liver sympathectomy.

We next asked whether this effect is liver-specific by performing selective hepatic sympathetic denervation before subjecting mice to SD. Like global sympathectomy, liver-specific denervation significantly attenuated SD-induced hepatic neutrophil infiltration and reduced neutrophil activation (Fig. 5c-d). Consistent with this, SD increased hepatic noradrenaline levels, whereas 6-OHDA treatment reduced UA levels (Supplementary Fig. 6a-c). Together, these results indicate that SD activates hepatic sympathetic signaling, which promotes UA synthesis and thereby drives neutrophil recruitment to the liver.

### Chemogenetic activation of hepatic sympathetic nerves enhances neutrophil influx

To establish a causal role for hepatic sympathetic signaling in neutrophil recruitment, we employed an intersectional chemogenetic strategy to selectively activate the sympathetic-hepatic circuit. We injected a dual-virus cocktail at multiple sites along the common hepatic nerve at the porta hepatis, before its branching in the liver. The first virus (AAV9.rTH.PI.Cre.SV40) drives Cre recombinase under the tyrosine hydroxylase (TH) promoter, restricting expression to catecholaminergic neurons, while the second (AAV9-hSyn-DIO-hM3D(Gq)-mCherry) expresses the excitatory DREADD hM3D(Gq) in a Cre-dependent manner. This strategy ensures that chemogenetic activation is confined exclusively to sympathetic nerve fibers innervating the liver. Four weeks after virus delivery, clozapine-N-oxide (CNO) was administered intraperitoneally to acutely activate hepatic sympathetic input, and neutrophil infiltration in the liver was assessed (Fig. 6a). Chemogenetic activation of hepatic sympathetic nerves resulted in a significant increase in hepatic neutrophil accumulation (Fig. 6b). These results provide direct evidence that activation of the hepatic sympathetic branch is sufficient to promote neutrophil recruitment, consistent with a model in which SD-induced sympathetic signaling enhances UA synthesis and thereby drives reversible neutrophil infiltration as a localized inflammatory response (Fig. 6c).

**Fig 6.**
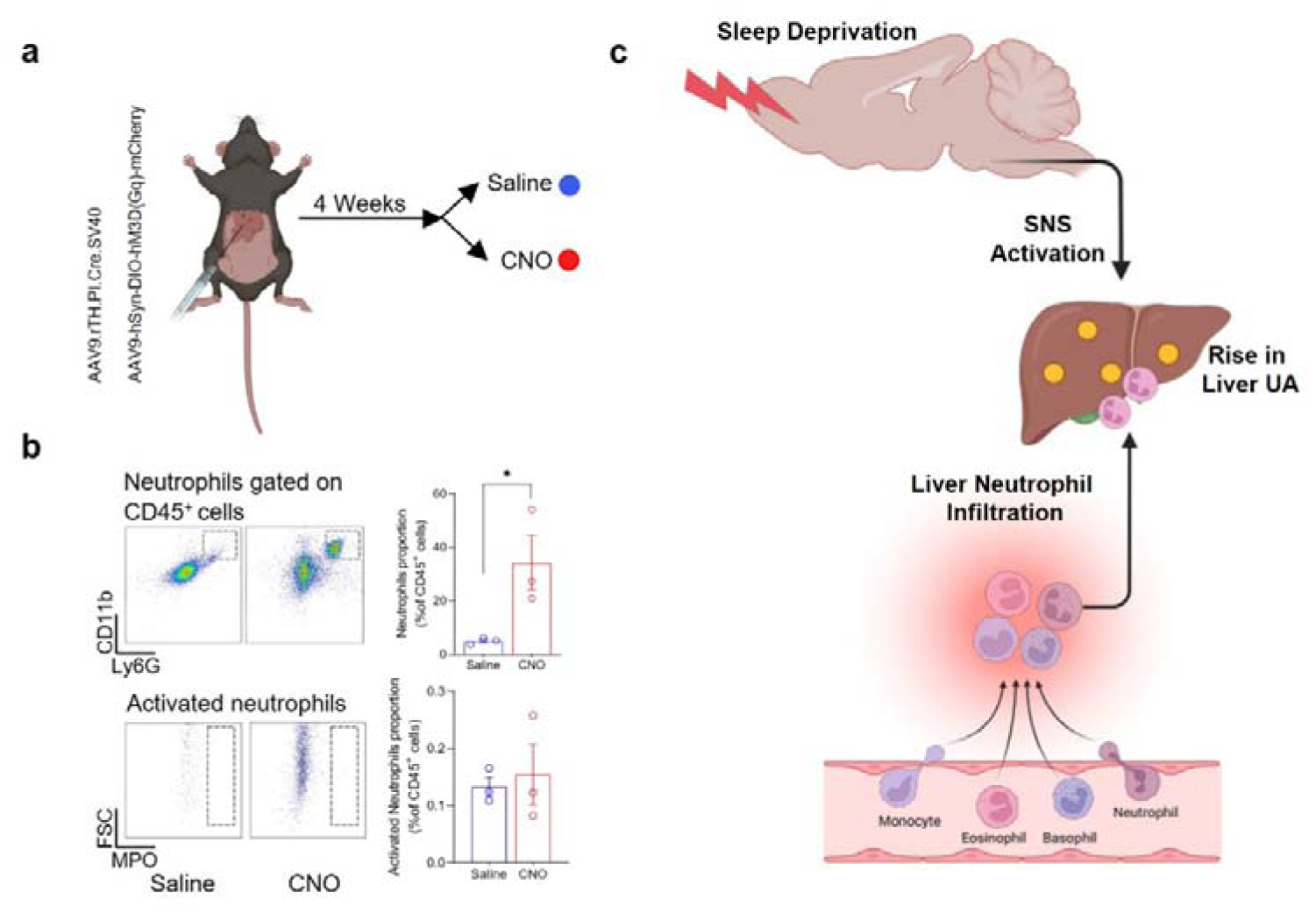
Chemogenetic activation of liver sympathetic tyrosine hydroxylase neurons enhances the hepatic neutrophil infiltrations. **a,** Schematic of chemogenetic activation of sympathetic tyrosine hydroxylase neurons achieved by AAV9.rTH.PI.Cre.SV40 and AAV9-hSyn-DIO-hM3D(Gq)-mCherry in the sympathetic nerve terminal entering to liver followed by CNO administration. b, Gating strategy and bar plots showing quantification for neutrophils (upper) and activated neutrophils (lower) in the liver of mice having sympathetic tyrosine hydroxylase neurons activated by CNO treatment. Data are mean□±□s.e.m. n□=□3 biological replicates. Unpaired t-test, two-tailed. *\*P* < 0.05. **c,** Schematic model of brain-sympathetic nervous system (SNS) mediated regulation of hepatic inflammation during sleep deprivation. Sleep deprivation activates SNS, increasing hepatic uric acid synthesis that triggers neutrophil accumulation in the liver.

## Discussion

Sleep profoundly influences metabolism and immune function, yet the mechanistic pathways through which sleep governs metaflammation remain poorly defined ^14, 27^. Here, we identify a sleep-dependent sympatho-metabolic-immune circuit in which acute sleep deprivation activates hepatic sympathetic signaling, enhances xanthine oxidase-dependent uric acid synthesis, and drives reversible neutrophil recruitment and sterile inflammation in the liver.

A key consideration in studying sleep loss is distinguishing its direct effects from stress-related confounds. We employed a validated 12-h acute SD protocol that eliminates ∼90% of sleep and produces robust slow-wave activity rebound during recovery, reflecting accumulated homeostatic sleep pressure ^16, 28^. Critically, this paradigm does not elevate plasma corticosterone, allowing us to attribute the observed metabolic and inflammatory responses to sleep loss itself rather than hypothalamic-pituitary-adrenal axis activation. By focusing on acute deprivation, we aimed to capture the earliest neuro-metabolic and immune events before compensatory or adaptive mechanisms obscure the primary signal.

Several converging lines of evidence point to the liver as the principal site of sleep loss-induced immune activation. Inflammatory mediators were broadly elevated in liver tissue but only selectively increased in plasma; hepatic UA levels rose markedly during SD, while circulating UA remained unchanged; and pharmacological inhibition of UA synthesis with allopurinol suppressed hepatic, but not systemic, inflammatory responses. The absence of elevated circulating UA, together with increased UA in liver lysates, implies that UA accumulates predominantly within or around hepatocytes during SD. This compartmentalization has important implications. UA is a well-established DAMP capable of triggering sterile inflammation ^21^, and its local accumulation within the liver, rather than systemic elevation, suggests that the inflammatory signal originates at the tissue level.

Consistent with this, MSU crystals are considerably more potent inducers of neutrophil chemotaxis than soluble UA ^20, 29^, and our *ex vivo* experiments confirm that MSU-stimulated hepatocytes robustly attract neutrophils. That hepatocyte-specific *Xdh* overexpression recapitulated SD-induced neutrophil infiltration *in vivo* further strengthens a causal link between intrahepatic UA elevation and neutrophil recruitment.

The sympathetic nervous system emerges as the critical upstream effector linking sleep loss to hepatic UA production. Prolonged wakefulness enhances sympathetic tone ^24, 25^, and sympathetic stimulation promotes hepatocyte UA synthesis ^26^. Our loss- and gain-of-function experiments converge on the conclusion that sympathectomy (both global and liver-specific) attenuated SD-induced neutrophil infiltration, while chemogenetic activation of hepatic sympathetic fibers was sufficient to drive neutrophil accumulation independently of sleep loss, establishing causality. A recent study reported that global sympathectomy does not alter the systemic rise in circulating neutrophils following SD ^14^. Our data are compatible with this finding but reveal a distinct, organ-selective role for sympathetic signaling in orchestrating local hepatic neutrophil influx, highlighting that systemic and tissue-specific neuro-immune programs operate in parallel during sleep loss.

What remains unresolved is how the brain senses accumulated sleep pressure and translates it into enhanced sympathetic output to the liver. The hypothalamic orexin system is a compelling candidate. Orexin neurons remain highly active during enforced wakefulness and project broadly to noradrenergic and sympathetic regulatory centres, including the locus coeruleus, rostral ventrolateral medulla, and paraventricular nucleus ^30, 31^. Chemogenetic activation of forebrain-PVN circuitry alone is sufficient to induce hepatic steatosis through sympathetic mechanisms ^32^, lending plausibility to a model in which SD-induced orexinergic activation engages sympathetic liver innervation to promote UA-dependent inflammation. Mapping this descending circuit in full will be an important future direction.

Our findings also intersect with the emerging picture of circadian immune regulation in the liver. Under homeostatic conditions, hepatic neutrophil surveillance follows a diurnal rhythm, peaking when sleep pressure is highest and declining at the onset of the active period, to coordinate daily metabolism in an inflammation-independent manner ^11, 33^. SD appears to override this temporal gating, driving pathological neutrophil accumulation during a phase when the liver would normally be immunologically quiescent. One interpretation is that enforced wakefulness co-opts innate immune programs alongside sympathetic arousal, mobilizing neutrophils as part of an anticipatory host-defence response. Such a response is evidently adaptive in the short term - it resolves completely with recovery sleep. However, when sleep loss becomes chronic, this same axis may become maladaptive, contributing to the enhanced haematopoiesis, sustained systemic inflammation, and cardiovascular and metabolic dysfunction documented in prolonged sleep disruption models ^34, 35^. The transition from reversible organ-specific inflammation to irreversible systemic disease represents a critical threshold that warrants further investigation, and the sympathetic-UA-neutrophil axis identified here offers a tractable entry point for therapeutic intervention aimed at mitigating metaflammation in the growing population affected by insufficient sleep.

## Acknowledgements

A.B.R. acknowledges funding from the Perelman School of Medicine, University of Pennsylvania, and the Institute for Translational Medicine and Therapeutics (ITMAT) at the University of Pennsylvania. This work was also supported by NIH DP1DK126167 and R35GM161590.

## Author contributions

P.K.J. and A.B.R. conceived the project and designed the experiments. P.K.J., U.K.V., and J.C. performed the experiments. P.K.J. analysed and interpreted data. A.B.R. supervised the entire study and secured funding. The manuscript was written by P.K.J. and A.B.R. with input from all authors. All authors agreed on the interpretation of data and approved the final version of the manuscript.

## AI Disclosure

Artificial Intelligence (AI) tools (GPT5.4 model) were used in the preparation of the written content of this manuscript only for editing and proof-reading (spell and grammar check).

## Methods

### Mice

All animal studies adhered to the approved University of Pennsylvania and ARRIVE guidelines. All animal experimental protocols received approval from the Institutional Animal Care and Use Committee (IACUC) at the Perelman School of Medicine, University of Pennsylvania. Wild-type 6-8 weeks old male C57BL/6J mice were obtained from Jackson Laboratories and allowed to acclimate for at least 2 weeks before the experiments. For sleep deprivation experiments, the mice were housed in automated sleep fragmentation chambers (Model #80391, Campden/Lafayette Instrument, Lafayette, IN, USA). At all times, the mice had ad libitum access to food and water under standard housing conditions, following a 12-h light: 12-h dark cycle.

### Cells

HEK293T cells were cultured in Dulbecco’s Modified Eagle’s Medium (DMEM; Thermo Fisher Scientific, 61965-026) containing 10% Fetal Bovine Serum (FBS) (F2442, Lot 21B456, Sigma-Aldrich) and 1% Penicillin Streptomycin (P/S) (15140122, Gibco) in an incubator with a humidified atmosphere at 37°C containing 5% CO_2_. Hepatocytes were isolated from the 8-10-week-old C57BL/6 mice by perfusing the mouse liver with 15 mL of reperfusion buffer (Hanks’ balanced salt solution, 0.5 mM EDTA), followed by 15 mL of collagenase digestion buffer (Dulbecco’s modified Eagle medium [DMEM] with low glucose, 15 mM HEPES, 0.5 mg/mL collagenase [C5138, Sigma], 0.1 mg/ml DNase I [DN25, Sigma], and 1% penicillin□streptomycin [P/S; Sigma]). Cells were released and passed through a 70-mm cell strainer and centrifuged at 50 g for 3 minutes. After washing with PBS, hepatocytes were resuspended and stored in DMEM with 10% FBS and 1% P/S. For primary hepatocyte culture experiments, 500,000 cells per well were plated in collagen-coated 24-well plates.

### Sleep deprivation and recovery sleep

Mice were divided into ad libitum Normal Sleep (NS), Sleep-Deprived (SD), and Recovery Sleep (RS) groups. The NS group was left undisturbed, while the SD group was sleep-deprived for 12 h [Zeitgeber Time 0-12 (ZT0-12)] using a device that applied a tactile stimulus with a horizontal bar sweeping just above the cage floor (bedding), based on our previous study ^16^. In addition to the sweeping bar, additional attempts were made to maintain wakefulness, mostly during the second 6□h of sleep deprivation by occasionally tapping on the cage or gently touching with a brush. The RS group was similarly sleep-deprived and allowed to recover for 24 h (ZT12-12). Post experiment, mice were sacrificed. A midline abdominal incision was made, and blood was collected by puncturing the inferior vena cava. Liver perfusion with PBS was performed as previously described ^36^, after which livers were harvested for preparation of single-cell suspensions and liver lysates for downstream analyses.

### Flow Cytometry to track liver-infiltrating neutrophils

Mice were perfused with 20 mL of PBS, and the entire liver was collected and dissociated using the gentleMACS™ Dissociator (MACS, 130-093-235) together with the Mouse Liver Dissociation Kit (130-105-807). The resulting tissue dissociates were passed through a 100-µm strainer to obtain a single-cell suspension. Cells were then washed and resuspended in PEP buffer (PBS, pH 7.2, containing 0.5% bovine serum albumin and 2 mM EDTA).

To identify neutrophils and activated neutrophils, cells were stained for 30 min at 4°C in the dark with the following antibodies: PerCP/Cyanine5.5 anti-mouse CD45 (BioLegend, 30-F11), APC anti-mouse/human CD11b (BioLegend, M1/70), PE anti-mouse Ly-6G (BioLegend, 1A8), and FITC anti-myeloperoxidase (Abcam, ab90812). Following staining, cells were fixed with 2% paraformaldehyde and analyzed by flow cytometry (Luminex 200). Data were processed using FlowJo software (v10.1).

### Complete Blood Count (CBC)

CBCs of blood were performed using the Abaxis hematology analyzer. The proportion of circulating lymphocytes, monocytes, and neutrophils was measured from total leukocyte counts.

### Hematoxylin and Eosin and Immunohistochemistry (IHC) Staining

Livers were fixed in 4% paraformaldehyde (PFA) overnight and subsequently embedded in paraffin. Sections of 10 µm thickness were prepared. After dewaxing and rehydration, nuclei were stained with hematoxylin and differentiated with acid alcohol, followed by eosin staining of the cytoplasm. Sections were then dehydrated and mounted with neutral resin. For IHC, deparaffinized and rehydrated sections were incubated with 3% hydrogen peroxide. After antigen retrieval, slices were blocked with 10% normal goat serum and were then incubated with primary antibody (Xanthine Oxidase, ab109235, 1:1000; Myeloperoxidase, ab208670, 1:1000). The Next day, sections were washed 3 times with PBS and incubated with secondary antibody for 1 hour. Imaging was performed using a Leica Biosystems Aperio AT2 Digital Pathology Scanner. Image J was used to quantify MPO spots and XO intensity.

### Plasma and liver cytokine assay

Cytokines in plasma and liver lysate were determined by mouse cytokines (Luminex), Millipore (MTH17MAG-47K), and mouse IL18 R&D 7625 70 following the manufacturer’s instructions. Each kit contained 96-well plates to determine different biomarkers; both plasma and liver lysate samples were appropriately diluted with dilution buffer for the assay according to the preliminary test.

### Enzyme-Linked Immunosorbent Assays (ELISA)

Both plasma and/or liver lysate levels of corticosterone (R&D system, KGE009), CPR (R&D MCRP00 5), MPO (ab275109), XO (ab102522), UA (ab65344), and noradrenaline (EA633/96) were determined by appropriate kits by following the manufacturer’s instructions.

### RNA extraction and Quantitative Reverse Transcription Polymerase Chain Reaction (qRT-PCR)

Total RNA was extracted from fresh frozen mouse cerebral cortex tissues using the Direct-zol RNA miniprep kit (R2052, Zymo Research). The extracted RNA was reverse-transcribed into cDNA using the High-Capacity cDNA Reverse Transcription Kit (4368814, Applied Biosystems) following the manufacturer’s instructions. qRT-PCR was performed using SYBR Green Master Mix for qPCR (A46110, ThermoFisher Scientific) and commercially available primers for Xdh (ID 22436), G6pc1 (ID 14377), Pck1 (ID 18534), Pparα (ID 19013), Pgc1α (ID 19017), Sirt1 (ID 93759), Nr1d1 (ID 217166), Ho-1 (ID 15368), IL-1β (ID 16176), Ccl2 (ID 20296), and Tlr4 (ID 21898). The RT-PCR reactions were run in duplicate for each target gene using a ViiA7 Real-Time PCR System (Applied Biosystem). Expression levels of target genes were determined using the delta-delta cycle threshold method to calculate fold changes, with normalization to beta actin expression levels.

### Targeted metabolomics assay and fluorometric assays

The upper right liver lobes were dissected and flash frozen. Lyophilized samples were used to map the metabolites involved in uric acid synthesis. Quantitation of metabolites was performed by liquid chromatography/mass spectrometry (two Agilent 1290 Infinity UHPLC/6495 triple quadrupole mass spectrometers). Fluorometric assays were performed to quantify Guanosine (Guanosine Assay Kit, MET-5149), Guanine (Guanine Assay Kit, MET-5147), and Xanthine/Hypoxanthine (Xanthine/Hypoxanthine Assay Kit, ab155900) in liver lysate.

### Drug treatments

Allopurinol (A8003, Sigma) at 15 mg/kg, prepared in 0.5% carboxymethyl cellulose (C5678, Sigma), was administered by oral gavage for 8 consecutive days, including the day of sleep deprivation, at ZT0. Clozapine N-oxide (CNO) dihydrochloride (3 mg/kg) and 6-hydroxydopamine (6-OHDA; Sigma-Aldrich, H4381; 120 mg/kg) were administered intraperitoneally. CNO was administered at ZT6, while 6-OHDA was administered at ZT0 for 5 consecutive days.

### Expression cassette design and AAV vector production

For Xdh/mCherry overexpression in the liver, we used pAAV.TBG.PI.Null.bGH (addgene #105536) as backbone and inserted Xdh or mCherry coding sequences after thyroxine binding globulin (TBG) promoter. For virus preparation, HEK293T cells were seeded in DMEM supplemented with 10% FBS and 1% Penicillin-Streptomycin the day before transfection. 60% confluent HEK293T cells were triple transfected with pAdDeltaF6, AAV2/8, and expression plasmid (pAAV-TGB-Xdh or pAAV-TGB-mCherry) at the same molarity using polyethyleneimine (43896, Thermo Scientific Chemicals). 90% of the culture medium was replaced by OptiPRO (12309019, Fisher Scientific) supplemented with 1% Penicillin Streptomycin 24 hours after the transfection. The cells were cultured in the medium for an additional 72 hours. Viruses were extracted using chloroform and precipitated in 50% PEG (Polyethylene glycol). AAVs were concentrated using AmiconUltra4 Centrifugal Filter Unit (UFC810024, Millipore) and stored at 4°C until use. The virus titer was measured by quantitative PCR on serially diluted samples using PowerUp SYBR Green Master Mix (Applied Biosystems) and primers against the ITRs, forward: 5’-GGAACCCCTAGTGATGGAGTT-3’, reverse: 5’-CGGCCTCAGTGAGCGA-3’, obtained from Sigma-Aldrich.

### Surgeries

For phenol-based hepatic sympathetic denervation, mice were anesthetized with isoflurane, and a midline abdominal skin and muscle incision was made to expose the hepatic artery, portal vein, and common bile duct. A sterile cotton-tipped applicator soaked in 10% phenol in ethyl alcohol was carefully applied to the surface of the hepatic artery and portal vein bundle. To minimize postoperative adhesion formation, the abdominal cavity was thoroughly rinsed with 0.9% saline. Sham surgeries were performed identically, except 0.9% saline was applied instead of phenol. For hepatic sympathetic activation, similar surgical procedures were performed. Instead of phenol application, the common hepatic branches at the porta hepatis, prior to intrahepatic branching were exposed to identify hepatic nerves, and DREADD viruses were microinjected at multiple sites. The abdomen was then closed using atraumatic sutures. Animals were allowed to recover for 10-12 days (phenol-denervation group) or 4 weeks (viral-injection group) before experimental use.

### Virus Administration

Mice received tail-vein injections of 200 µL (2 × 10^11^ gc) of AAV2/8-TBG-Xdh or AAV2/8-TBG-mCherry. Seven days after injection, animals were sacrificed, and livers were collected to assess hepatic neutrophil infiltration. For liver sympathetic activation, a viral cocktail (10^13^ vg/mL) consisting of AAV9.rTH.PI.Cre.SV40 and AAV9-hSyn-DIO-hM3D(Gq)-mCherry were delivered to hepatic nerves. Four weeks later, animals were sacrificed, and livers were harvested to evaluate neutrophil infiltration.

### MSU crystal synthesis

MSU crystals were prepared by recrystallization from UA as previously described ^37^. Briefly, 1 g of UA and 6 mL of 1 mol/L NaOH were dissolved in 194 mL of deionized water, and the mixture was heated until fully solubilized. The pH was then adjusted to 7.2 using 1 mol/L HCl, and the solution was allowed to crystallize at room temperature overnight. The resulting precipitate was collected by centrifugation at 200 × g for 5 min and washed three times with 75% ethanol. Crystals were dried at 60°C overnight, sterilized, resuspended in fresh sterile DPBS, and stored at 4°C until use.

### Isolation of WBCs from peripheral blood

Mice were euthanized, and peripheral blood was collected from the inferior vena cava using EDTA-flushed syringes to prevent coagulation. RBCs were lysed by adding RBC lysis buffer (0.8% ammonium chloride, 4.3 mM potassium bicarbonate, and 0.1mM EDTA) at a volume ten times that of the collected blood and incubating the mixture at room temperature for 10 min. Subsequently, three volumes of culture medium (DMEM) were added, and samples were gently inverted to mix before centrifugation at 500 × g for 5 min at 4°C. The lysis and centrifugation steps were repeated until RBC removal was complete and a visible white cell pellet was obtained. The final cell pellet was resuspended in culture medium. Cell numbers were determined using an automated cell counter (Countess II Automated Cell Counter, Cat. #AMQAX1000).

### Transwell assay

For transwell migration assays, 200,000 adult mouse WBCs were seeded into 5-µm pore inserts (CellQart, Sterlitech; 24-well PET inserts) and placed into 24-well plates containing one of the following in the lower chamber: culture medium alone, medium with MSU, supernatant from hepatocytes treated with MSU, or hepatocytes treated with MSU. Transwells were incubated at 37°C for 2 h to allow migration. Cells that migrated to the lower wells were collected after 2 h. Cells were stained to identify the proportion of migrated neutrophils with PerCP/Cyanine5.5 anti-mouse CD45, APC anti-mouse/human CD11b, PE anti-mouse Ly-6G. Following staining, the cells were fixed with 2% paraformaldehyde and analyzed by flow cytometry (Luminex 200). Data were processed using FlowJo software (v10.1).

## Figure Legends

**Supple. Fig. 1.**
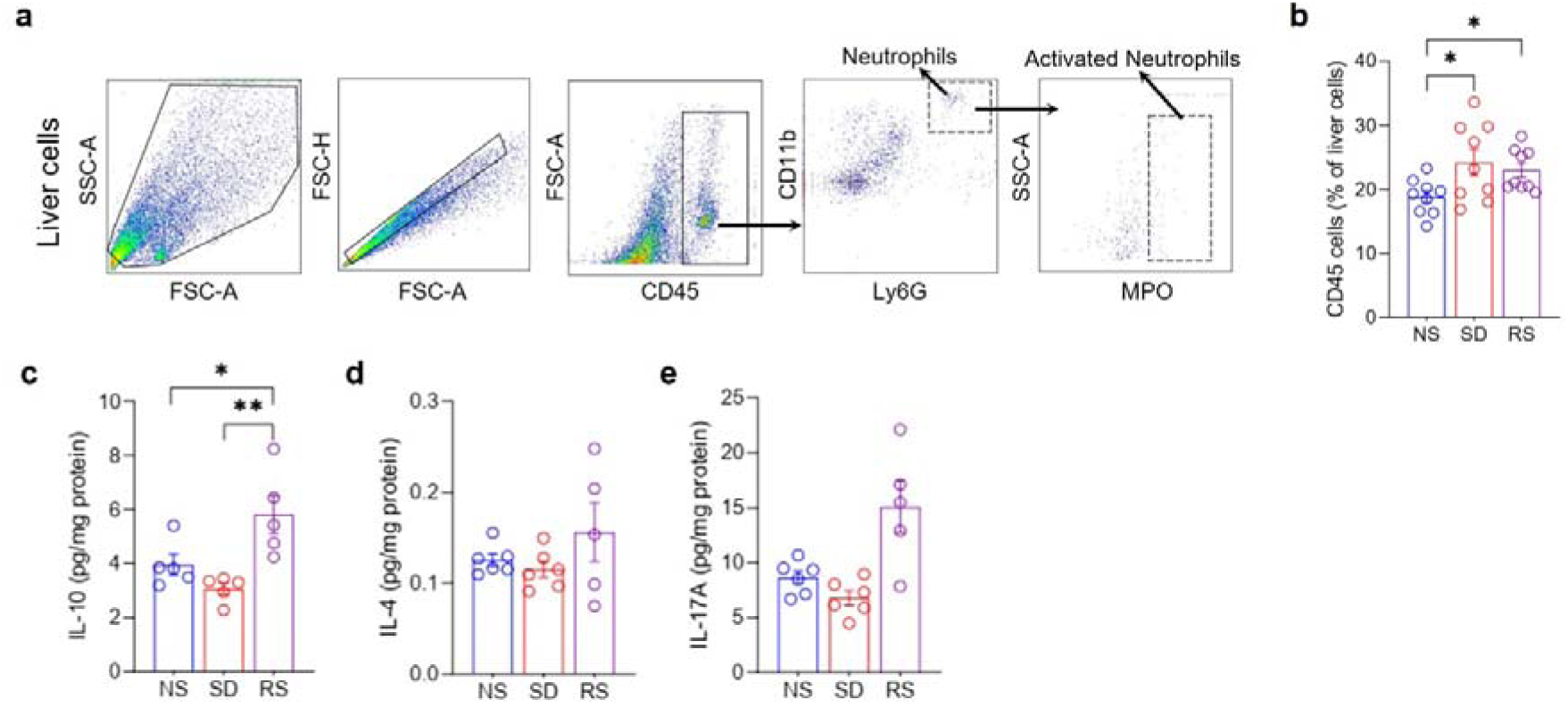
Sleep regulates hepatic CD45 cell levels and inflammatory mediators. **a,** Gating strategy for neutrophils (CD45+CD11b+Ly6G+) and activated neutrophils (CD45+CD11b+Ly6G+MPO+) in liver cell suspension. **b-e,** Bar plots showing alterations of liver CD45 cells (b), IL-10 (c), IL-4 (d), and IL-17A (e) across sleep treatments. Data are mean□±□s.e.m. n□=□5-9 biological replicates. One-way ANOVA followed by post-hoc Fisher’s Least Significant Difference (LSD) test. *\*P* < 0.05, and *\*\*P* < 0.01. NS normal sleep, SD sleep deprived, RS recovery sleep.

**Supple. Fig 2.**
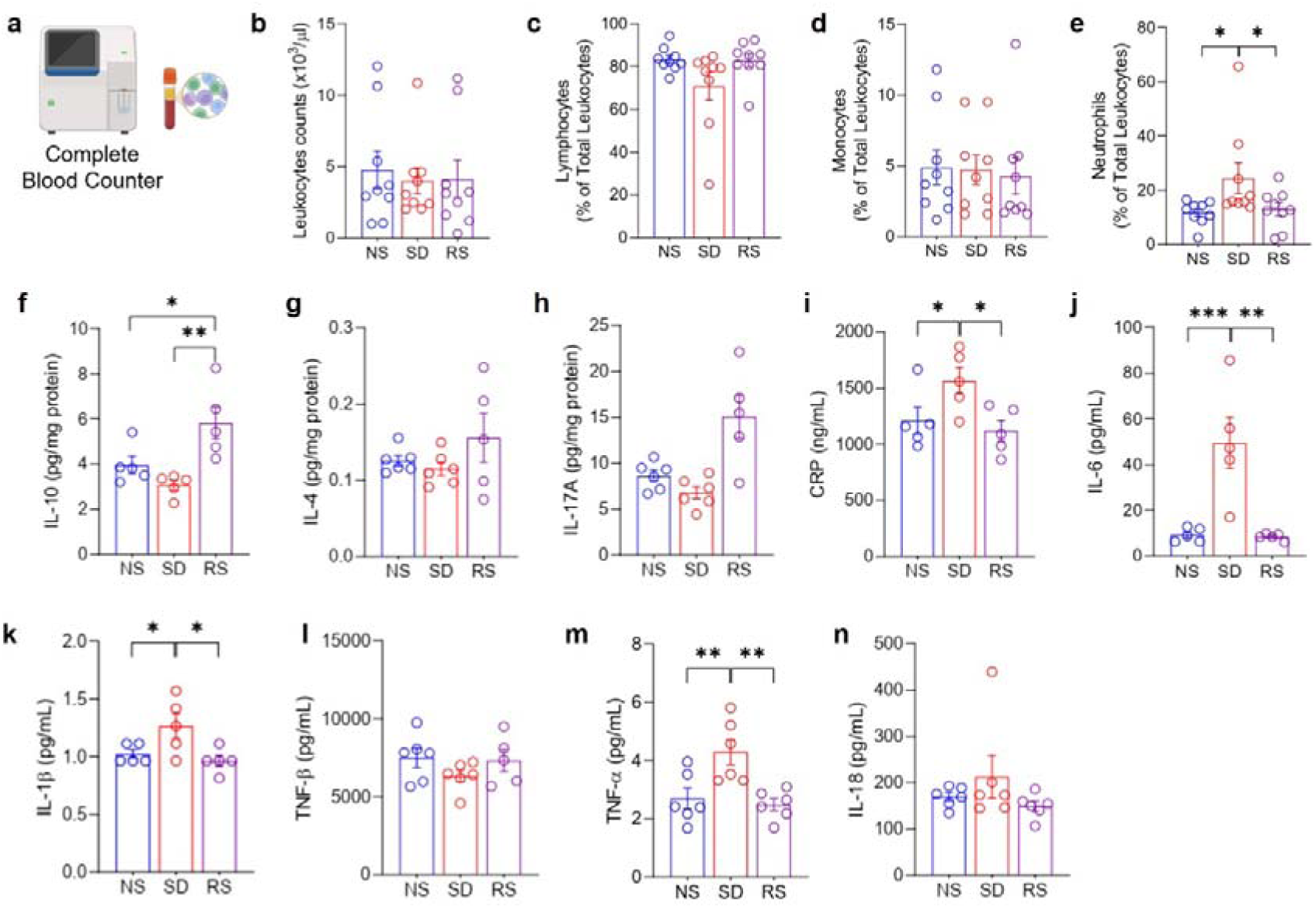
Sleep treatments alter the circulating neutrophils and inflammatory mediators. **a,** Schematic for complete blood counts enumeration. **b-e,** Bar plots showing total leukocytes (b) and the proportion of lymphocytes (c), monocytes (d), and neutrophils (e) in the mice subjected to sleep treatments. **f-n,** Plasma levels of IL-10 (f), IL-4 (g), IL-17A (h), CRP (i), IL-6 (j), IL-1β (k), TNF-β (l), TNF-α (m), and IL-18 (n) in the mice subjected to sleep treatments. Data are mean□±□s.e.m. n□=□3-9 biological replicates. One-way ANOVA followed by post-hoc Fisher’s Least Significant Difference (LSD) test. *\*P* < 0.05, *\*\*P* < 0.01, and *\*\*\*P* < 0.001. NS normal sleep, SD sleep deprived, RS recovery sleep.

**Supple. Fig 3.**
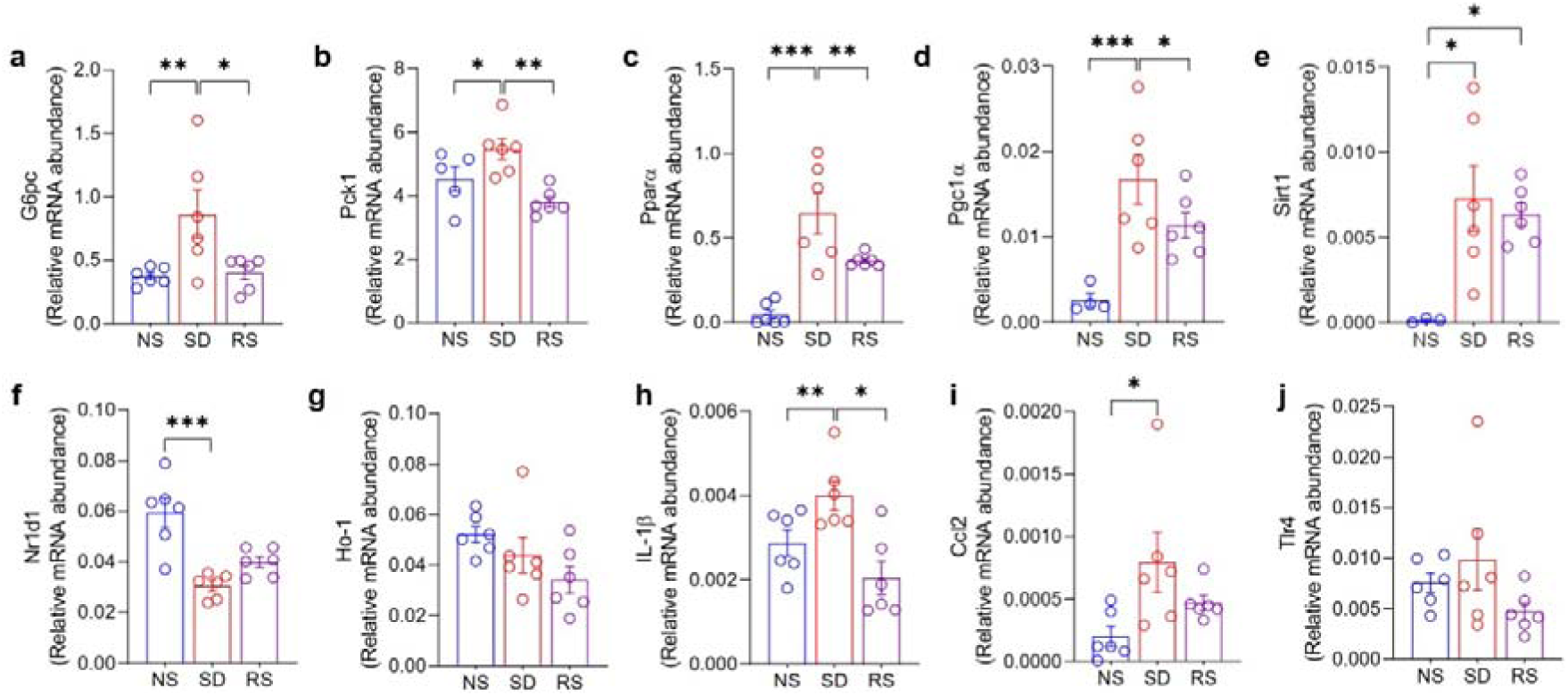
Sleep treatments alter hepatic gene expression associated with metabolism and inflammation. **a-j,** Bar plots showing mRNA expression of *G6pc* (a), *Pck1* (b), *Ppar*α (c), *Pgc1*α (d), *Sirt1* (e), *Nr1d1* (f), *Ho-1* (g), *IL-1*β (h), *Ccl2* (i), and *Tlr4* (j) normalized to *Actb*. Data are mean□±□s.e.m. n□=□3-6 biological replicates. One-way ANOVA followed by post-hoc Fisher’s Least Significant Difference (LSD) test. \**P* < 0.05, \*\**P* < 0.01, and \*\*\**P* < 0.001. NS normal sleep, SD sleep deprived, RS recovery sleep.

**Supple. Fig 4.**
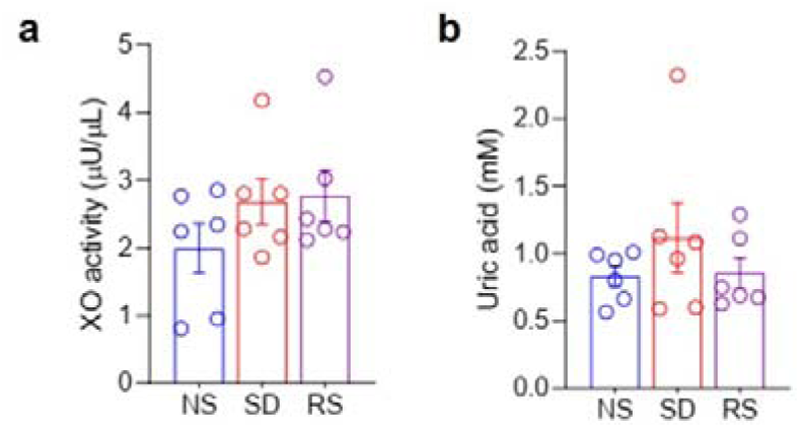
Sleep deprivation enhances liver corticosterone. **a-d,** Bar plots showing corticosterone levels from the liver lysate (a), plasma corticosterone (b), XO activity (c), and uric acid (d) of animals subjected to different sleep treatments. Data are mean□±□s.e.m. n□=□6 biological replicates. One-way ANOVA followed by post-hoc Fisher’s Least Significant Difference (LSD) test. *\*P* < 0.05. NS normal sleep, SD sleep deprived, RS recovery sleep.

**Supple. Fig 5.**
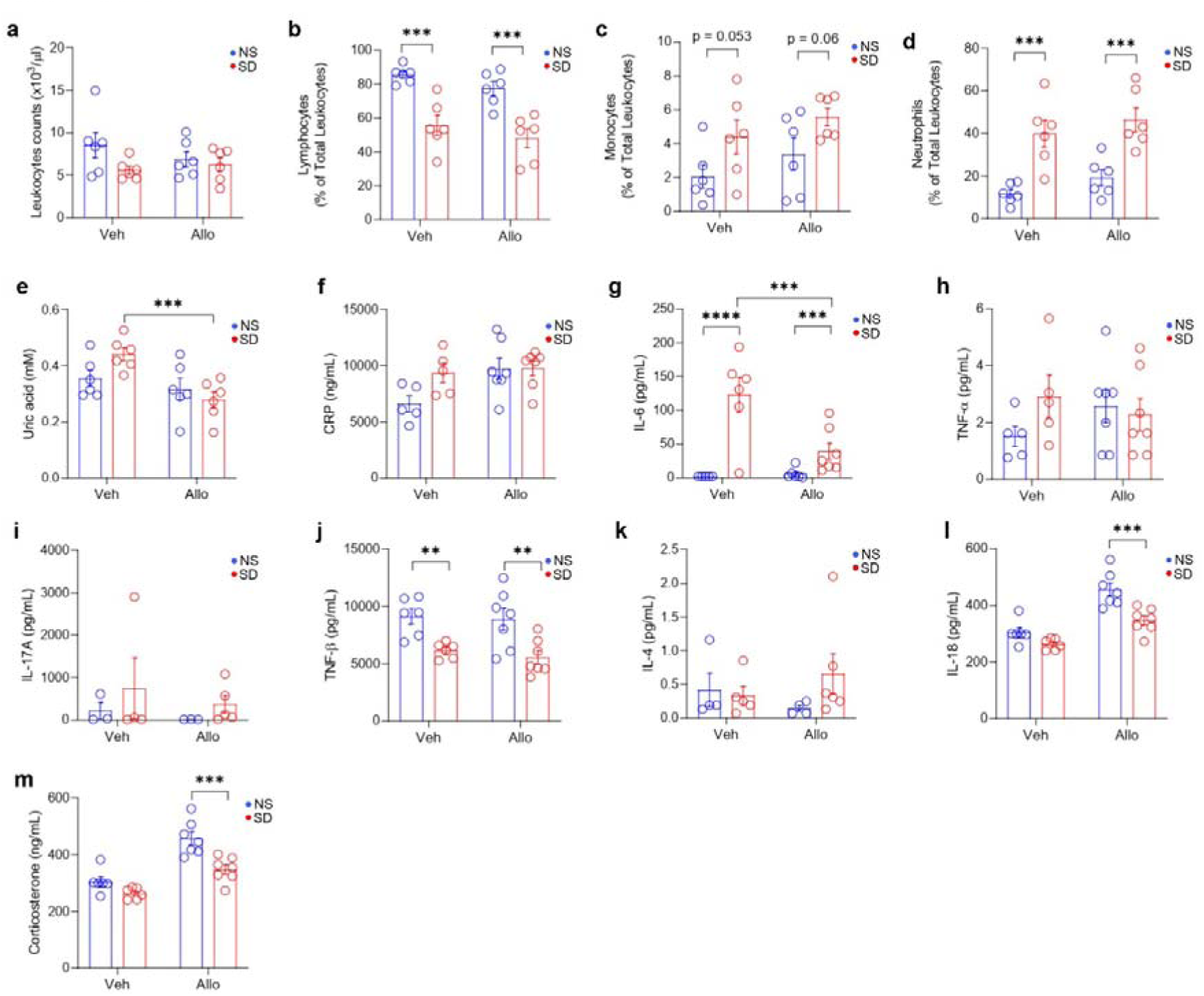
Allopurinol remains ineffective in lowering circulating neutrophils and plasma levels of inflammatory mediators induced by sleep deprivation. **a-d,** Bar plots showing total leukocytes (a) and the proportion of lymphocytes (b), monocytes (c), and neutrophils (d) in the mice treated with Allo/Veh followed by sleep treatments. **e-m,** Plasma levels of uric acid (e), CPR (f), IL-6 (g), TNF-α (h), IL-17A (i), TNF-β (j), IL-4 (k), IL-18 (l), and corticosterone (m). Data are mean□±□s.e.m. n□=□3-7 biological replicates. Two-way ANOVA followed by post-hoc Fisher’s Least Significant Difference (LSD) test. *\*P* < 0.05, *\*\*P* < 0.01, and *\*\*\*P* < 0.001. NS normal sleep, SD sleep deprived, Veh vehicle, Allo allopurinol.

**Supple. Fig 6.**
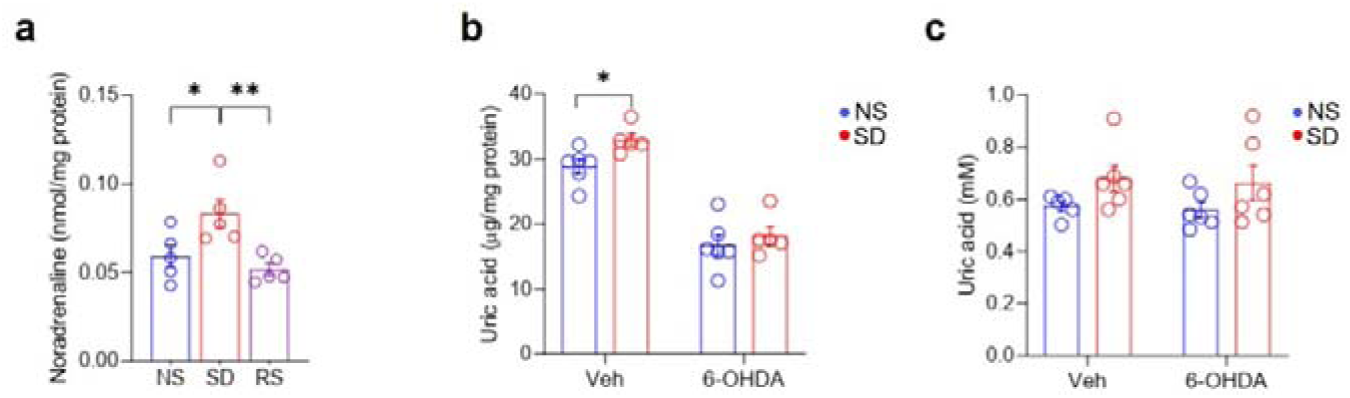
Sleep deprivation increases hepatic noradrenaline, and 6-OHDA reduces hepatic urate levels. **a-c,** Bar plots showing alterations of noradrenaline across sleep treatments (a), uric acid in the liver (a), and plasma (c) in chemically sympathectomized mice subjected to sleep deprivation. Data are mean□±□s.e.m. n□=□5-6 biological replicates. One-way (a) and two-way (b-c) ANOVA followed by post-hoc Fisher’s Least Significant Difference (LSD) test. *\*P* < 0.05 and *\*\*\*P* < 0.001. NS normal sleep, SD sleep deprived, RS recovery sleep.

